# Computational Analysis of Dynamic Allostery and Control in the SARS-CoV-2 Main Protease

**DOI:** 10.1101/2020.05.21.105965

**Authors:** Igors Dubanevics, Tom C.B. McLeish

## Abstract

The COVID-19 pandemic caused by the novel coronavirus SARS-CoV-2 has generated a global pandemic and no vaccine or antiviral drugs exist at the moment of writing. An attractive coronavirus drug target is the main protease (M^pro^, also known as 3CL^pro^) because of its vital role in the viral cycle. A significant body of work has been focused on finding inhibitors which bind and block the active site of the main protease, but little has been done to address potential non-competitive inhibition which targets regions beyond the active site, partly because the fundamental biophysics of such allosteric control is still poorly understood. In this work, we construct an Elastic Network Model (ENM) of the SARS-CoV-2 M^pro^ homodimer protein and analyse the dynamics and thermodynamics of the main protease’s ENM. We found a rich and heterogeneous dynamical structure in the correlated motions, including allosterically correlated motions between the homodimeric protease’s active sites. Exhaustive 1-point and 2-point mutation scans of the ENM and their effect on fluctuation free energies confirm previously experimentally identified bioactive residues, but also suggest several new candidate regions that are distant from the active site for control of the protease function. Our results suggest new dynamically-driven control regions as possible candidates for non-competitive inhibiting binding sites in the protease, which may assist the development of current fragment-based binding screens. The results also provide new insight into the protein physics of fluctuation allostery and its underpinning dynamical structure.

## 1. Introduction

In 2019, a rapidly spreading disease named COVID-19 caused by the novel coronavirus SARS-CoV-2, has since generated a global pandemic. Preventive measures have been taken by a majority of countries, but no vaccine or anti-viral drugs exist, at the time of writing, although candidates are under trial. In the longer term, the identification of all potential inhibitor sites at all points of the viral life-cycle is of interest. Here we focus on the low-frequency dynamical structure of the virus’ main protease, an important molecular machine in the viral cycle, and identify critical residues in its allosteric control. The work is informative for inhibitor design by identifying control regions of the protein that are distant from, rather than proximal to, its active sites. Allosteric mechanisms for distant control of binding and activation fall into two main classes: those which invoke significant conformational change (the original scenario of Monod, Wyman and Changeaux (1), and mechanisms that invoke the modification of thermal (entropic) fluctuations about a fixed, mean conformation (2–5). Such ’fluctuation allostery’ recruits mostly global, low-frequency modes of internal protein motion, which are well-captured by correspondingly coarse-grained mechanical representations of the protein (6, 7). One effective tool at this level is the Elastic Network Model (ENM) (8). The ENM resolves protein structure at the level of alpha-carbon sites only, which are represented as nodes connected by harmonic springs within a fixed cut-off radius from each other. Local point mutation can be modelled by changing the moduli of springs attached to the corresponding residue, and effector-binding by the addition of nodes and local harmonic potentials. The most significant contributions to the correlated dynamics of distant residues, and to the entropy of fluctuation come from global modes, whose ENM approximation allows straightforward calculation. This approach was successfully used to identify candidate control residues whose mutation may control allostery of effector binding in the homodimer transcription factor CAP (9). This study, and others, have shown that, while the huge reduction in the number of degrees of freedom that the ENM constitutes, does not capture the quantitative values of free energies, or their changes on mutation that are seen in experiment, it can identify the qualitative nature of the thermal dynamics of a protein. Furthermore, its coarse-graining can determine which residues present as candidates for allosteric control through mutation. The method, and the open software (’DDPT’) (10) used in the previous study on allosteric homodimers is deployed here in a similar way (see Methods section) to a coarse-grained ENM model of the SARS-CoV-2 Main Protease.

### A. The SARS-CoV-2 Main Protease Protein

At the time of this study (July 19, 2020) more than 37,000 papers have been published in relation to the virus (See COVID-19 Primer). However, work is still in progress to identify biological and molecular pathways the virus takes. Fortunately, significant research has already been directed to very similar coronavirus - SARS-CoV, first identified in 2003. The SARS-CoV and SARS-CoV-2 genetic sequences are almost 80% identical (11). Both viruses encode the main protease (M^pro^), also known as the 3C-like protease (3CL^pro^). In its active form M^pro^ is a two protomer homodimer with one active site per the homodimer chain (12). Although M^pro^ is a relatively compact protein (less than 310 residues per chain), it plays a vital role in the viral cycle of both coronaviruses: it divides polyproteins expressed from the viral mRNA into its functional non-structural units (13). This functional role makes SARS-CoV-2 M^pro^ an appealing target for drug design. The major research effort to date has been focused on the competitive inhibition of SARS-CoV-2 M^pro^, i.e. by directly targeting the active site with molecules that competitively bind to the active site “pockets” (11, 14–16). A significant body of work has been recently published investigating inhibitors for the SARS-CoV-2 main protease via virtual screening and MD simulations (17–24). In 2011 it was found that N214A mutation dynamically inactivates SARS-CoV M^pro^ (25).

The same research group later characterised another SARS-CoV M^pro^ mutation, S284-T285-I286/A, which dynamically enhances the protease catalytic activity more than three-fold (26). In SARS-CoV-2 M^pro^ two of those amino acids (T285 and I286) are changed to T285A and I286L with respect to SARS-CoV M^pro^. However only a little enhancement in catalytic activity is observed (11). This delicate potential control region will appear below in our analysis (see section B.2.). Due to the high sequence conservation between both coronaviruses, and 96% amino acid sequence identity between the main proteases, SARS-CoV-2 M^pro^ might posses similar allosteric features. Furthermore, in another study, researchers found the the root mean square deviation (RMSD) of 0.53Å for apo (ligand-free) forms of two corona viruses’ main proteases (PDB accession 2BX4 and 6Y2E) (11). These evidence that the N214A mutation operates through a fluctuation allostery mechanism and structural similarities between two proteases motivates the analysis of the coarse-grained dynamic structure of SARS-CoV-2 M^pro^ reported here. We apply the ENM techniques of (9) to this purpose, looking in particular to identify non-active, yet allosteric, sites for non-competitive inhibition.

## 2. Simulations, Results and Discussions

No crystallographic structure of the SARS-CoV-2 M^pro*^ active form with a polyprotein is available to date. Only empty (apo) structures or structures with synthetic ligands/substrates attached to the active site available. Therefore, the ENM study reported here used a recent crystallographic structure (PDB accession 6LU7 (14)) with competitively inhibited active site to calculate fluctuation free energies and consequent allosteric energies and their modification under mutation. The inhibited (with α-ketoamide inhibitors, referred as “holo2”) and ligand-free (apo) structures of the protein are almost identical shown in figure 1. Resolutions for the structures shown are 2.16Å and 1.75Å respectively, while RMSD between them is under 1.5 Å for Cα atoms. Evidently, very little structural change happens upon the inhibitor binding. These findings further support the hypothesis of dynamically driven allosteric control of SARS-CoV-2 M^pro^, and provide a structure (6LU7) on which to base an ENM construction (**SI, Sec. A**).

**Fig. 1.**
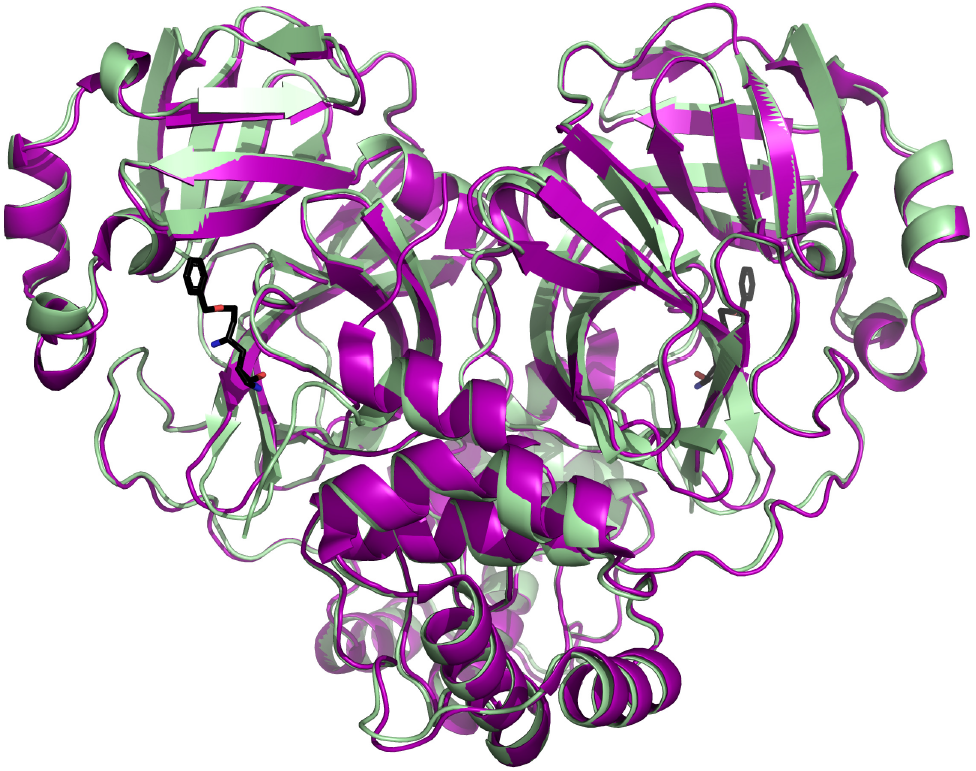
Inhibited (pale-green) and ligand-free (purple) SARS-CoV-2 M^pro^ crystallographic structures superimposed using the Combinatorial Extension (CE) algorithm in PyMOL (Schrödinger). The RMSD between two structures is 1.48Å. Both proteins are shown as secondary structure cartoons while an α-ketoamide inhibitor is shown in licorice representation. PDB accessions are 6LU7 and 6Y2E, respectively.

The resulting ENM of M^pro^ is shown in figure 2. It takes Cα node masses as the whole residue mass, and uses a cut-off distance for harmonic connecting springs of 8Å, based on comparison of mode structures with full Molecular Dynamics simulations in previous work on Catabolite Activator Protein (9, 27)(**SI, Sec. B**). Balancing the requirements of: (i) sufficient spatial resolution of dynamics; (ii) requirements not to include unphysically small-scale structure; (iii) acceptable convergence of thermodynamic calculations; (vi) compatibility with the previous studies (28) leads to the choice of summing the first real 25 modes in SARS-CoV-2M^pro^ ENM calculations (**SI, Sec. C**).

**Fig. 2.**
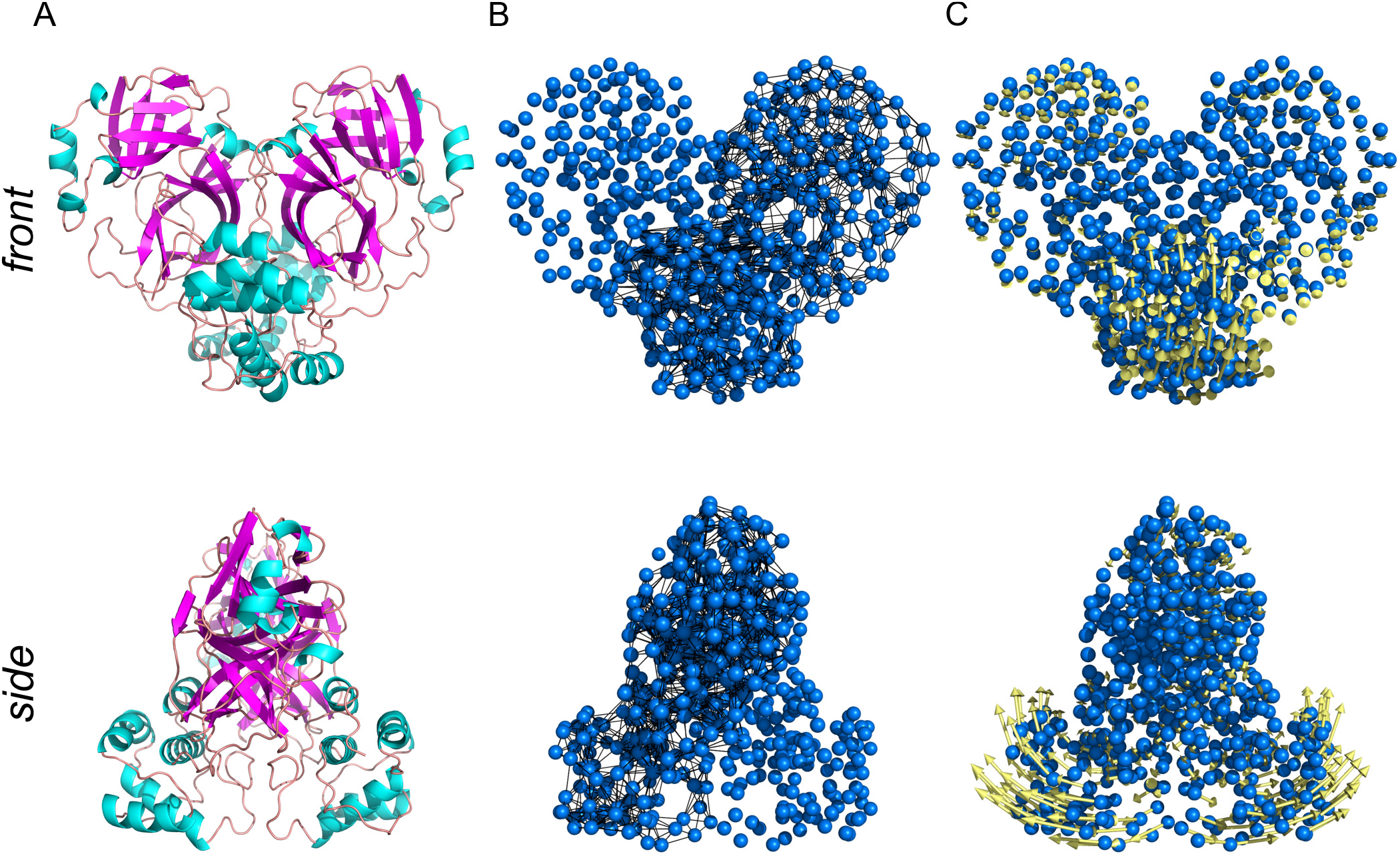
Constructing ENM of SARS-CoV-2 M^pro^ step-by-step. (A), SARS-CoV-2 M^pro^ secondary structure cartoon (B), Elastic model of M^pro^ generated with PyANM package in PyMOL. Cα atoms are shown in blue; while node-connecting springs (black) are shown only for one chain for comparison. (C), The first real vibrational mode eigenvectors (yellow) visualisation. For clarity, displacement vectors are scaled 5 times.

### A. Residue-residue dynamic cross-correlation map

The first quantity of interest is the map of residue-residue cross-chain dynamic correlations, which indicates for each residue on the protein those other residues whose motion correlates with its own (**Eq. 1**). This gives both a detailed summary map of the homodimer dynamical structure, and is also significant thermodynamically since the same elastic communication drives both correlations and allosteric control (4). The dynamic cross-correlation of motion map for the all residues in the ENM apo (ligand-free), holo1 (only one active site at chain A occupied) and holo2 (both active sites occupied) structures are shown in figure 3. We discuss the dynamic features of each structure in the following:

**Fig. 3.**
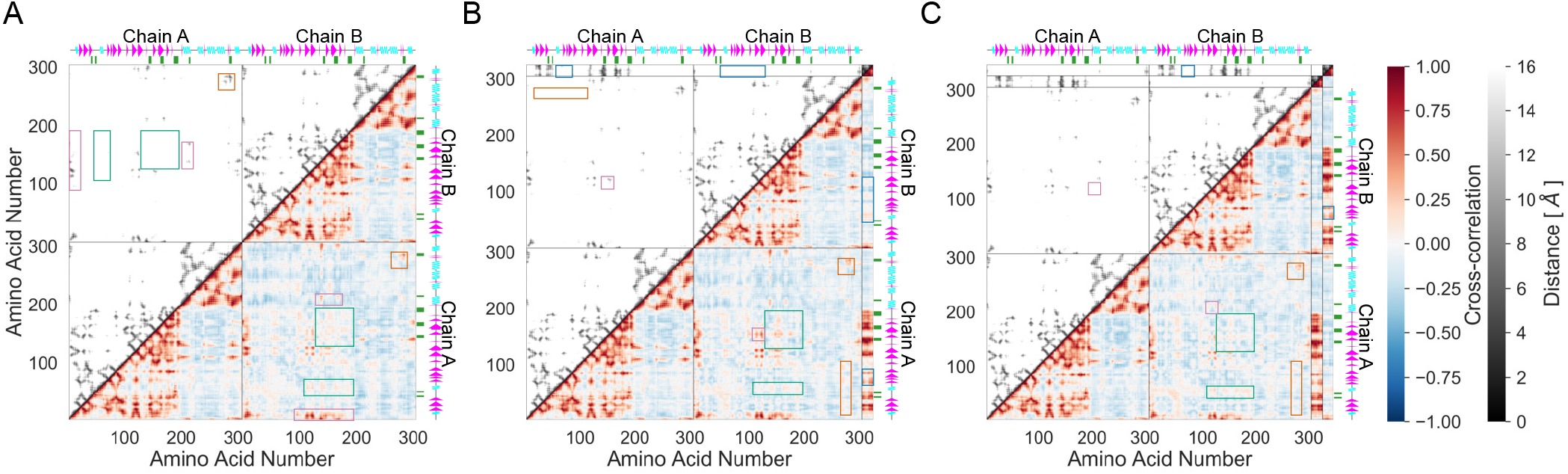
The cross correlation of the motion for the first real 25 modes and distance between residues(Cα nodes) as shown in 2-dimensional residue space for (left) apo, (*middle*) holo1 and (*right*) holo2 forms of 6LU7 ENM. The first colour scale show the extent of cross correlation, with a cross correlation of 1 (red) indicating perfectly correlated motion, −1 (blue) showing perfectly anti-correlated motion and 0 (white) no correlation. The second colour scale (black to white) depicts the Euclidean distance between two Cα nodes in the Cartesian space in 0-16Å range. The secondary structure of M^pro^ is indicated along the residue axes, with cyan waves indicating alpha helices, and magenta triangles indicating beta sheets. The green ticks on the axis indicate the location of the biologically active residues (**Tab. 1**). The cross correlation matrix was calculated using only the Cα atoms for the protein and all heavy atoms for ligand (α-ketoamide inhibitor).

#### A.1. apo

The N-terminus of each chain positively correlates with residues adjacent to the active site (res 100-200) on the other chain. This is due to physical proximity rather than allostery (**Fig. 3A**, wide purple rectangles in lower right and upper left quadrants). Significantly, the dynamics of active sites on both chains positively correlate, and allosterically, with each other (**Fig. 3A**, green square and rectangle in lower right quadrant). These regions are spatially far away: we can not see them at the corresponding location on the distance map (**Fig. 3A**, green square and rectangle in upper left quadrant). The T201-N214 alpha helix (which contains the experimentally-sensitive N214) on one chain dynamically anti-correlates with H41 on the opposite chain. The same helix, from residue 201 to 213, also anti-correlates with C145 on the opposite chain; while surprisingly (since its mutation is effective in allosteric control) N214 shows no correlation with the catalytically vital C145 at all. We observe strong positive correlation between this helix and two regions forming the active site pocket: residues K137-N142 (loop) and E166-H172 (*β*-turn) (**Fig. 3A**, narrow purple rectangles in lower right and upper left quadrants). However, this correlation can partially be accounted for by spatial proximity. S284-L285-A286 residues (henceforth SLA) on one monomer show positive cross-chain dynamic coupling of motion with the identical residues on the other, in this case through spatial proximity (**Fig. 3A**, orange squares in lower right and upper left quadrants), but a somewhat smaller positive correlation spans residues 275 to 306 (N terminus) on both chains with respect to SLA. This effect suggests strong SLA coupling to a large fraction of the protein domain not containing the active site. Furthermore, SLA positively correlates with the T201-N214 alpha helix, which contains the experimentally determined dynamically allosteric residue N214. Beyond the helix, G215 and D216 show slightly greater correlation with SLA. Finally, SLA negatively correlate with the active site’s catalytic dyad, H41 and C145, and residues around it (**Fig. 3A**, not shown). Thus, we observe both dynamic correlation at a distance, as well as that due to immediate spatial proximity, supporting previous findings regarding SARS-CoV M^pro^ dynamically allosteric inactivation.

#### A.2. holol

The positively correlated dynamics between the active sites are strongly enhanced by an addition of the first ligand, especially in regard to two beta sheets (G146-I152 and V157-L167) and a beta turn between them (**Fig. 3B**, green square in the lower right quadrant). Interestingly, and by contrast, the region displaying positive correlation around residues 50-70 in the apo form is decreased in the holo1 structure (**Fig. 3B**, green rectangle in the lower right quadrant). In the holo-1 form, the structural symmetry of the apo form is broken, permitting the asymmetric magnitude of correlation between chains A and B (**Fig. 3B**), across the diagonal of the lower right quadrant). The biologically active residues (green ticks) show up in the ligand’s correlation with (host) chain A. However, four other regions, not cited in the literature to date, also show strong dynamic correlation with the ligand. Two of them (res 17-32 and 120-131) can be vividly observed as spatially proximal to the ligand from the corresponding distance map. Two other regions exhibiting positive cross-correlation are, however, distant from the ligand (**Fig. 3B**, blue rectangles in lower right and upper left quadrants). These regions span residues 67-75 and 77-91, respectively, and include two beta sheets and a beta turn in each case. The previously reported dynamically allosteric residue 214 (chain B) correlates positively with ligand on chain A; potentially due to spatial proximity. The closest ligand’s residue to N214 is 11.2Å away. Moreover, the ligand shows positive correlation at distance with the beta sheets on the opposite chain (**Fig. 3B**, blue rectangles in upper right quadrant). The fact that the ligand’s motion positively correlates with beta sheets at residues 67-75 and 77-91 on both chains suggests strong chain-ligand coupling around residues 67-91 region on both homodimer chains.

The dynamically allosteric SLA and the region around dynamically couples to the same residues on the opposite chain (**Fig. 3B**, orange square in lower right quadrant). The structural *C_2_* symmetry breaking upon the ligand binding decouples motion of residues which are far away form the active site (**Fig. 3B**, orange square in lower right quadrant). Thus these residues can engage in a collective motion driven by spatial proximity to their neighbours. Not seen on apo cross-correlation map, the second half of the N261-N274 alpha helix on chain B appears to correlate positively with four distant residue groups in 15-100 region on chain A (**Fig. 3B**, orange rectangle in upper left and lower right quadrants). Moreover, addition of the ligand enhanced produced an ’H’ shape correlation (on the cross-correlation map) between four beta sheets, two on each chain: G146-I152 and V157-L167 on chain A, G109-Y118 and S121-R131 on chain B (**Fig. 3B**, purple square in lower right quadrant). This group motion across chains is partially caused by two neighbouring beta turns sticking out of active site protein domains (**Fig. 3B**, purple square in upper left quadrant).

#### A.3. holo2

When a second ligand is added, the strong correlation between active sites present in holo1 is diminished (**Fig. 3C**, green square in lower right quadrant); while previously lowered correlation reaches the apo form level (green rectangle in the same quadrant and figure). The region around SLA and the N261-N274 alpha helix, which shows strong dynamical coupling with the residue groups in interval 15-100 of the holo1 form, is reduced on binding the second effector to its previous apo level correlation (**Fig. 3C**, orange square and rectangle in lower right quadrant). The newly added ligand on chain B shows the same correlation-at-distance with the L66-A70 and V73-L75 beta sheets, excluding the beta turn in between them (**Fig. 3B**, blue rectangles in upper right quadrant). Nevertheless, the cross chain coupling between active site and the alpha helix (T201-N214) is further increased. The correlation is split into three distinctive regions in locally ’wedge-like’ shapes on the map, two of which are around active site. These structures were also seen in the CAP cross-correlation map (9), and indicate the two contributing beta strands acting as a local hinge region. The third region is located around a beta sheet at S121-R131, which is not bound to the ligand (**Fig. 3C**, purple squares in lower right and upper left quadrants). The cross-correlation displays an interesting structure along the helix from residue N201 to T214: the third region is split into three zones: positive, negative and positive correlation. This sign change reminds 3-rd harmonic of a standing wave with two nodal points, whereas in 3D those are nodal planes. The negative correlation region is absorbed by the two positive regions as we reach residue N214. Unsurprisingly, the two ligands exhibit identical correlations with their host and opposite chains. Noteworthy, each ligand has dynamics correlating positively with the termini of the opposite chain, arising principally from the spatial proximity of C and N termini to the opposite chain’s active site.

### B. Mutation scans for thermodynamic control

The ENM calculations were extended to calculations of fluctuation (entropic) free energies (**Eq. 2**) of various modelled wild type and mutated state of SARS-CoV-2 M^pro^. ’Point mutations’ are modelled in these ENM calculations, as in (9) by softening or stiffening all the harmonic springs attached to each residue in a complete scan of equally spaced spring moduli with centre at 1.00 in range from 0.25 to 4.00. Figure 4A reports the free energy changes (*G_mut_ − G_wt_*)/|*G_wt_*| induced in the entire homodimer when this point mutation is done. Figure 4C reports the effect of the same mutational scan on the allosteric free energy *K*_2_/*K*_1_ (**Eq. 3**) of binding between the two active sites.

**Fig. 4.**
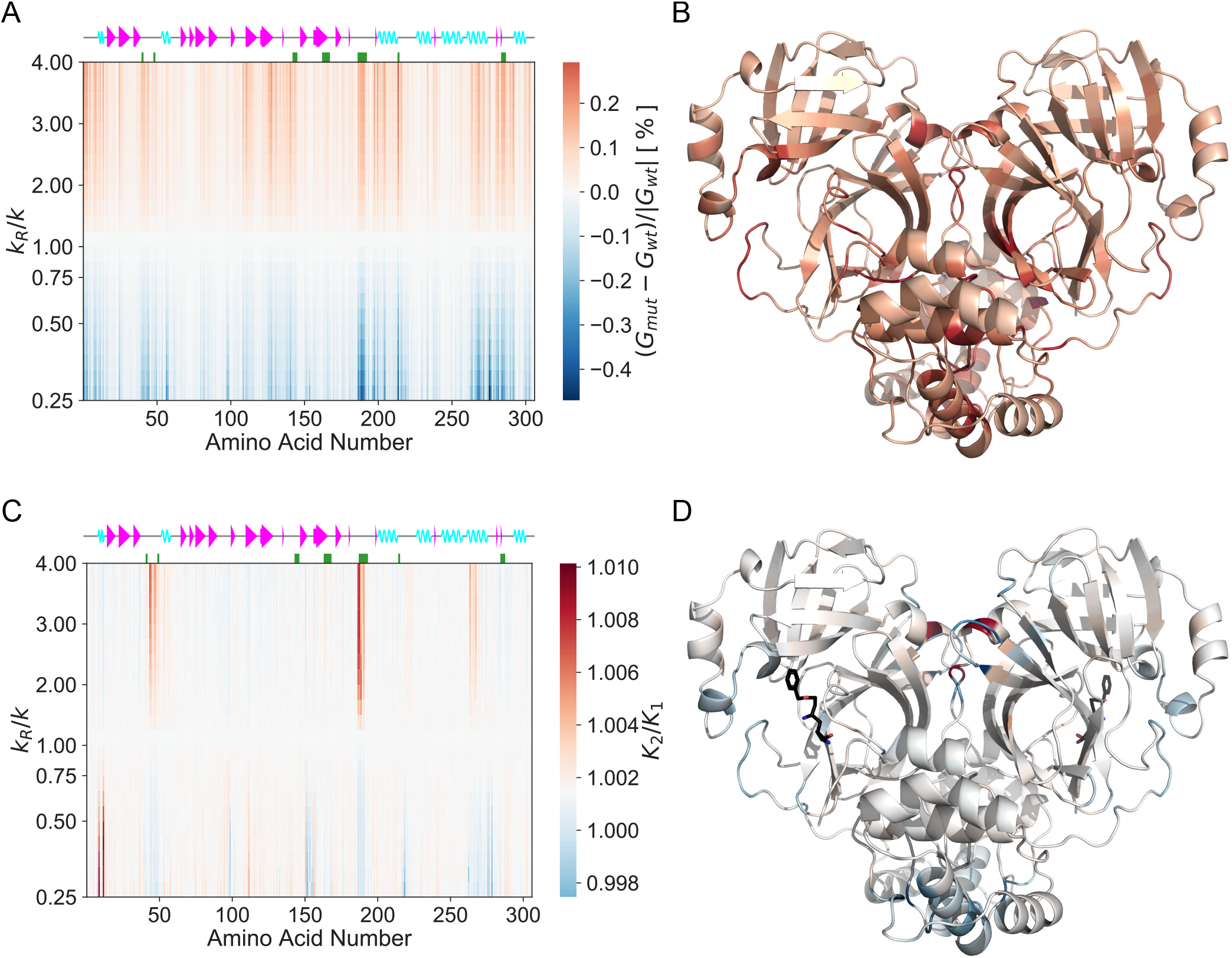
Mutation scan maps for thermodynamic control of M^pro^ calculated from the ENM over the first real 25 fluctuation modes. (A) A map for the fluctuation free energy change. The map plots the relative change in free energy to the wild type (*(G_mut_ – G_wt_*)/|*G_wt_*|) due to the dimensionless change in the spring constant (*k_R_*/*k*) for the mutated residue with the amino acid number shown. White corresponds to values of free energy predicted by the wild-type ENM. Red corresponds to an increase in (*G_mut_ – G_wt_*)/|*G_wt_*| (decreased value of *G_mut_* comparing to *G_wt_*), whereas blue corresponds to a decrease in ((*G_mut_ – G_wt_*)/|*G_wt_*|) (increased value of *G_mut_* comparing to *G_wt_*). (B) The map for the vibrational free energy change plotted in real space onto the wild-type M^pro^ homodimer structure at *k_R_*/*k*=4.00. (C) A map for the global control space of allostery in M^pro^. The map plots the change in cooperativity coefficient (*K*_2_/*K*_1_) due to the dimensionless change in the spring constant (*k_R_*/*k*) for the mutated residue with the amino acid number shown. White corresponds to values of *K*_2_/*K*_1_ predicted by the wild-type ENM. Red corresponds to an increase in *K*_2_/*K*_1_ (stronger negative cooperativity), whereas blue corresponds to a decrease in *K*_2_/*K*_1_ (weaker negative cooperativity or positive cooperativity). (D) The global map plotted in real space onto the wild-type M^pro^ homodimer structure at *k_R_*/*k*=0.25.

#### B.1. Free energy mutation scan on apo structure

All experimentally identified active sites (besides res 163-167) appear on the mutation scan of the 6LU7 apo structure, a somewhat remarkable result considering that no ligand is present to emphasise the spatial nature of the active sites. They seem dynamically pre-disposed to dynamic allosteric communication, in agreement with the cross-correlation map (**Fig. 3**). Both termini display mutation peaks due to their spatial proximity to the active sites. Additionally, a very sharp peak is seen around residues 187-192 where a free loop forming the active site is located. The seven new regions seen on the cross-correlation map (res 17-32, 67-75, 77-91, 97-98, 120-131, 201-214 and 261-274) form distinctive peaks as well on the mutation scan.

Furthermore, the experimentally identified residue N214 is signposted by these calculations: its computational mutation generates the largest fluctuation free energy change upon spring stiffening of 0.29 % at *k_R_*/*k*=4.00. The largest negative free energy change value of −0.47 % is produced upon spring relaxation of M276 at *k_R_*/*k*=0.25. These two points define the amplitude of the colour bar. Although the catalytically paramount residue C145 is not as sharp as other peaks, it appears with greater strength in a higher mode summation (**SI, Fig. S3A**). The map is mostly qualitatively anti-symmetric around middle line (wild-type). However, the quantitative behaviour of three regions worthy of attention: the region around residue 50, before 100, at residue 150, as well as the sharpest region around residues 187-192. Relaxation of stiffness at those points cause larger energy change than stiffening. A very narrow region, not identified on the cross-correlation map, at residues 97-98, preceding a small beta sheet at res Y101-V104, appears sharply when local interaction strengths are relaxed.

A new broad region of strong sensitivity to mutation appears on this map at residues 261-293, which includes an alpha helix at V261-N274. This helix is located on the surface of the protein far from the active site. This region also contains SLA which appear as sharp lines in figure 4A; and is especially responsive to spring constant change at L285: a free-energy change of 0.19 % at *k_R_*/*k*=4.00 and −0.29 % at *k_R_*/*k*=0.25. L285 is in the middle residue of the triad affecting two of its neighbours; furthermore, L285 on one chain is in the closest contact with its counterpart on the other chain (5.3 Å).

We also note 7 residues which are located on the homodimer chains’ interface (K5, P9, K12, E14, M276, I281 and S284), recalling that in the CAP homodimer, residues located on the interface were critical in allosteric regulation (9). Especially responsive is E14 located on the very first alpha helix.

#### B.2. Allosteric free energy mutation scan

The first result from the ENM calculation of the allosteric free energy for binding-site occupation is that *K*_2_/*K*_1_ ≈ 1 for the wild-type M^pro^ ENM. Therefore, this ENM model (over 25 softest modes) is non-cooperative. Nevertheless, we can identify regions that are sensitive to even slight change in local stiffness which again are around biologically active areas. All previously-marked active regions show-up to some extent; especially vivid is the region around catalytic residue H41 and, as already appeared in the apo mutation scan, the loop around the active site (res 187-192). Residue N214 shows very weak allosteric control in this scan. The local environment around N214 is mainly hydrophobic (**SI, Fig. S4**). Therefore, the experimentally-reported N214A mutation corresponds to a local structural stiffening (asparagine (N) is hydrophilic while alanine (A) is hydrophobic). This is indeed the region of parameter space (*k_R_*/*k*>1) where this mutation displays a weak effect in the ENM model, but a strong response upon relaxation.

In SARS-CoV-2 M^pro^ residues T285 and I286 are replaced by L285 and A286 with respect to SARS-CoV M^pro^. Purely from the perspective of hydrophobicity of residue and environment, the former mutation would correspond to *k_R_*/*k*>1, while the latter emulates *k_R_*/*k*<1. However, no exact comparison with experimental data can be made as there is no data on how S284-L285-A286/A simultaneous mutation affects SARS-CoV-2 M^pro^ catalytic activity. Without data on single mutations within the SLA region, we have no direct experimental verification of the single S284A mutation which in our ENM corresponds to *k_R_*/*k*>1 for similar reasons as for N214A mutation (**SI, Fig. S4**). We see a decrease in cooperativity for S284 spring stiffening, while for spring relaxation the ENM’s cooperativity increases.

### C. 2-point mutational scans

It is of interest to explore the cooperative effect of two-point mutations in models of fluctuation allostery, as previous work has indicated that double mutations may combine non-linearly in control of the allosteric landscape of proteins (27). This numerical scan explores cases where mutations are made not only on one of the single homodimeric chains: experimentally this is a possible, but not a trivial, task. However it can reveal contribution of each chain alone to fluctuation and allostery of the dimeric composite structure. The discussion of this sections refers to results presented in the 2-point scans of 6LU7 ENM in figure 5. In order to present the response to all double-mutations on a single 2D plot, the change in spring stiffness is not scanned; rather, just two constant spring changes of 0.25 and 4.00 are considered. Spring-stiffening 2-point mutational scans (**Fig. 5A,C**), *k_R_*/*k*=4.00, models the effect of small molecule/ligand binding to the mutated residues (and would also model mutations such as N214A (**SI, Fig. S4**)); while *k_R_*/*k*=0.25 map looks at the opposite extreme to the stiffening case, which would model mutations that weaken local bonding (**Fig. 5B,D**).

**Fig. 5.**
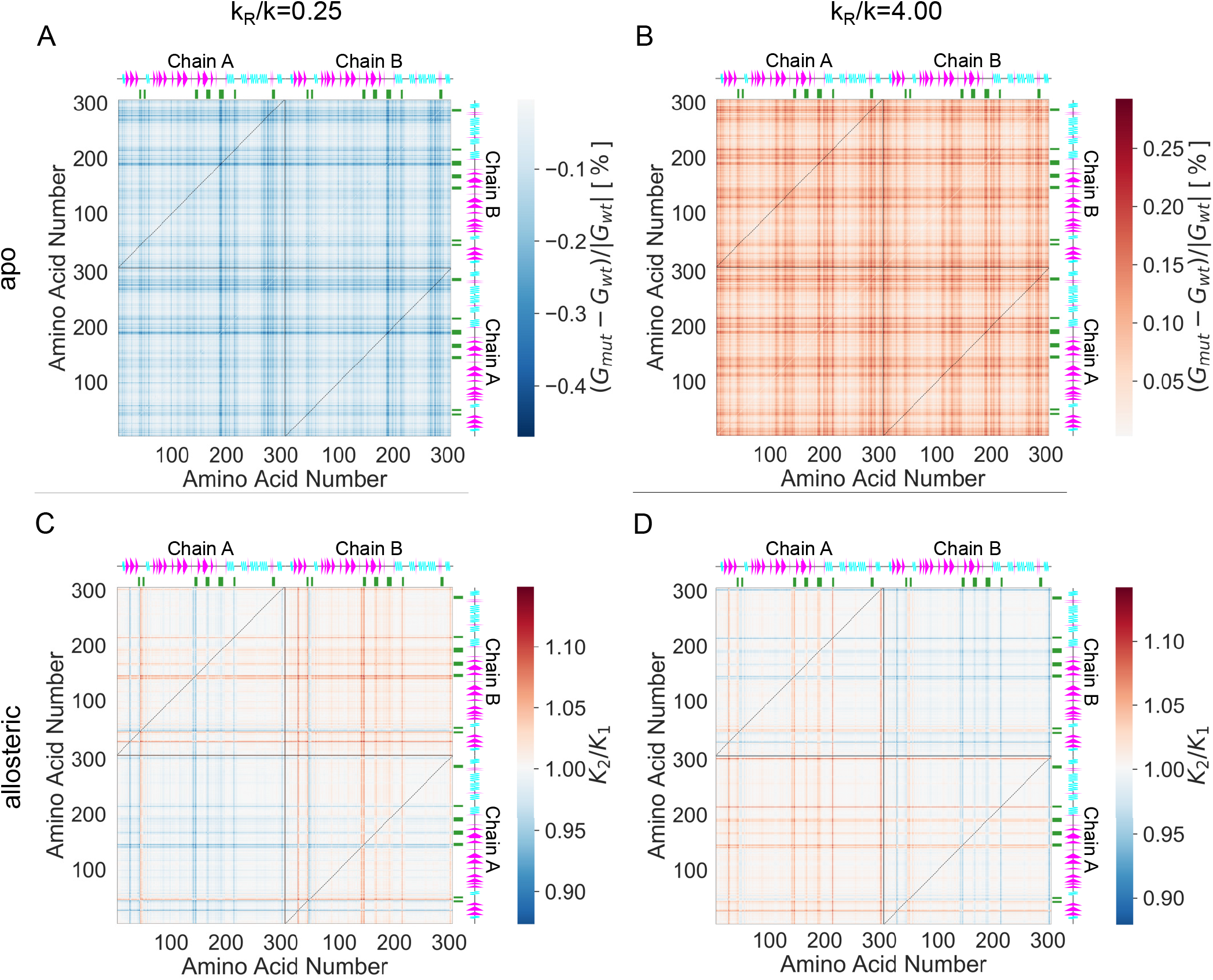
2-point mutational maps for 6LU7 ENM with all possible pairwise combinations of residue mutations with equal spring constant change *k_R_*/*k* equal 0.25 and 4.00 over the first real 25 fluctuation modes. (A,B) 2-point mutational maps for 6LU7 ENM with all possible pairwise combinations of residue mutations with equal spring constant change (A) *k_R_*/*k*=4.00 (spring stiffening) and (B) *k_R_*/*k*=0.25 (spring relaxation). (C,D) A map for the 2D global control space of allostery in M^pro^ for (C) *k_R_*/*k*=4.00 and (D) *k_R_*/*k*=0.25. Black solid lines separate two homodimer chains, while dashed lines represent 1-point mutational scan results for the given spring constant change.

#### C.1. Free energy 2-point mutation scan on apo structure

The first measure, as in the single-point scans, is the difference in total free energy of the apo structure. As on the 1-point map for apo 6LU7 structure strong lines are observed (**Fig. 5A,B**), but spring relaxation resolves fewer biologically active residues than spring stiffening. In figure 5A only residues around H41, a loop region forming active site at D187-A191 and N214 show up as solid lines. Whilst, stiffening (**Fig. 5B**) resolves all bioactive residues except H163-L167 (beta sheet forming the active site pocket) with with an additional region around a small beta sheet (Y101-V104), alpha helices (T201-N214 and V261-N274) and loop region adjacent to the latter helix. As in case of the 1-point map (**Fig. 4A**) M276 (−0.47 %) and N214 (0.29 %) define maximum absolute response upon relaxation and stiffening of these residues on both chains, respectively (**Fig. 5A,B**). In both cases the SLA region show moderate fluctuation free energy change. We conclude that stiffening is a better choice for resolving critical residues in fluctuation free energy control. Note that, in the case of this protein, the 2-point mutations combine approximately linearly: the effect of the first mutation (vertical lines) is not strongly affected by the second mutation. Nevertheless, response to relaxation and stiffening qualitatively different plots.

#### C.2. Allosteric free energy 2-point mutation scan

Finally, the effect of all double mutations upon spring relaxation and stiffening on the allosteric free energy between the two active sites was calculated (**Fig. 5C,D**). While the 2-point apo maps show qualitatively different behaviour for relaxation and stiffening, the allosteric free energy 2-point mutation maps are qualitatively exactly the same with an inverted allosteric free energy change sign. A new strong control site that did not appear on the 2-point apo maps (**Fig. 5A,B**) is found at the beginning of the second beta-sheet (T25-L32) on each chain (**Fig. 5C,D**). Additionally, S1 and G302 exhibit strong allosteric control due to spatial closeness to the active site and, thus, the ligand. The apparent increased cooperativity for both mutations (in most cases) on chain A, and the same pattern of decreased cooperativity on chain B, is due to the formal symmetry breaking through choice of the first binding site at chain A. Mutation of residues around H41 are the exception, and have an opposite effect on allostery to all others. The dashed lines represent the 1-point allo scan in figure 4 for corresponding *k_R_*/*k* values. In that scan *K*_2_/*K*_1_ values range from 0.998 to 1.010, while for these 2-point mutational scans the range is slightly increased.

This calculation draws attention to an additional advantage of the 2-point scans: while N214 did not exhibit allosteric control on the 1-point map (**Fig. 4C**) over 25 fluctuation modes, it appears as a strong line in all quadrants on the 2-point maps. However, when the lines intersect in the cross-chain quadrant *K*_2_/*K*_1_ reaches almost wild-type value (corresponding to making exactly the same change on both monomers). This effect explains why we do not observe allosteric control of N214 on the 1-point mutational map: evidently, when the bold lines intersect (identical mutation or binding on both domains) allosteric effects interfere destructively. Thus, in the 1-point allo scan for 6LU7 ENM amplitude of *K*_2_/*K*_1_ is lower than in the 2-point mutational allosteric scan or hardly shows up. The SLA region appears neither on the 1-point nor 2-point global maps for dynamic regulation of allostery in spite of its presence on the free-energy apo scans. This absence of control indicates the limited coupling between SLA and the given ligand (α-ketoamide inhibitor).

## 3. Discussion: what does ENM tell us about SARS-CoV-2 Main Protease?

The ENM analysis reinforces previous findings in application to other proteins, that in the SARS-CoV-2 M^pro^ as well, local harmonic potentials within the equilibrium protein structure, but without mean structural change can identify already known biologically active sites. Furthermore, there is no need to have holo forms of the protein to locate those active sites, whose correlated dynamics are already clear in the apo form. Calculations of those sites where total free energies are sensitive to mutations converge well with the limit of the sum over normal modes. The convergence of calculations of control of the allosteric free energy itself is more subject to noise, being a difference-quantity, but sufficiently to identify strong candidates for control regions (**SI, Fig. S3**).

The analysis shows that SARS-CoV-2 M^pro^ possesses a rich dynamical structure that supports several long-distance allosteric effects through thermal excitation of global normal modes. In particular the motions in the vicinity of two active sites are correlated within the first 25 non-trivial normal modes, especially in the singly-bound dimer. Although, at the level of ENM calculations, this does not lead to cooperativity in the WT structure, it does render the protein susceptible to the introduction of cooperativity by mutation.

Our methodology is further supported by the ENM dynamics sensitivity to residue 214 and 284-286 mutation which has been previously experimentally verified to dynamically control SARS-CoV M^pro^. The ENM calculations have identified new sites whose local thermal dynamics dynamically correlate with those of the active sites, and which also appear on global maps for allosteric control by single or double mutations. The new candidate control regions are summarised in table 1. In particular, residues around the beta sheets (Y101-V104, G109-Y118 and S121-R131) and the alpha helices (T201-N214 and V261-N274) are novel, and distant from the active site (**SI, Fig. S5**). The position of these residues suggest them as possible candidates for non-competitive inhibiting binding sites. We also draw attention to eight residues located on M^pro^ interface surface (**Tab. 1**) as a potential dynamically allosteric control residues.

**Table 1.** SARS-CoV-2 Main Protease key information used in this study: PDB ID; experimentally identified bioactive residues of SARS-CoV M^pro^ reported in literature; active regions which we identified in this study, distant from the active site for SARS-CoV-2 M^pro^ (SI, Section D).

| Protein | PDB ID | Experimentally Identified Bioactive Residues | Computationally Identified Distant Control Residues (this study) |
| --- | --- | --- | --- |
| SARS-CoV-2 Main Protease | 6LU7 | 41, 49, 143-5, 163-7, 187-192 (18)<br>214, 284-6 (25, 26) | 5, 9, 14, 109, 111, 127, 197-8,<br>200, 205, 214, 264, 268-9,<br>272, 276-7, 281-2, 284-6, 292 |

Computational studies such as this, therefore, accompany and support concurrent experimental programs of scanning for small-molecule binding candidates to the protein in question. We note that several candidate molecules, identified in a very recent large crystallographic fragment screen against SARS-CoV-2 M^pro^ (29), bind to regions suggested as dynamically sensitive control candidates in this study.

The ENM model employed was specific for the given inhibitor. Other ligands might, of course, show different behaviour in the corresponding holo structures and display other “hot-spots”, however, the appearance of active regions, and their coupling, in the apo structure suggests that there are general properties that emerge from the global elastic structure of the protein. As well as providing specific information on the SARS-CoV-2 M^pro^ structure of the calculations reported there, the findings of this study also contribute to the large programme of research on fluctuation-induced allostery without conformational change. In particular the general question of the focusing of dynamic correlations between distant (so candidate allosteric) sites is solved in a highly specific way by this structure. It also constitutes a system for which double mutations contribute in a predominantly linear addition, in contrast to findings with other allosteric homodimers. This pattern includes the cancellation phenomenon we identified in the case of some single point mutations made identically and simultaneously in both monomers, whose cancellation in the 1-d scans can mask their potential sensitivity as target sites. Finally, the appearance of control regions on the exterior surface of proteins, with obvious pharmacological application, generates other general questions in the biophysics of fluctuation elasticity in globular proteins.

### Materials and Methods

Normal Mode Analysis (NMA) of ENM describes protein motions around equilibrium and can be used to calculate the partition function for large scale harmonic thermal fluctuations in protein structure, including those responsible for allostery (30). Two main approximations of NMA are:

- The structure fluctuates about at local energy minimum. Consequently no other structures beyond the given equilibrium can be explored.
- The force field everywhere arises from sums over ENM harmonic

The whole NMA method can be reduced to three steps:

1. Construct mass-weighted Hessian for a system. For a protein ENM the system consists of the co-ordinates of the C-alpha atoms (*N*) for each residue from the corresponding PDB structure.
2. Diagonalise the mass-weighted Hessian to find eigenvectors and eigenvalues of the normal modes.
3. Calculate the partition function (and so free energy) from the product over the normal mode harmonic oscillations.

The diagonalisation of the 3*N* × 3*N* mass-weighted Hessian matrix is written as

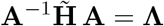

where 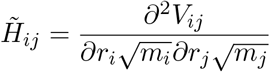: the potential energy function *V*; distance between nodes *r*; node masses *m*. The eigenvectors of the mass-weighted Hessian matrix, columns of **A**, are the normal mode eigenvectors **a**.

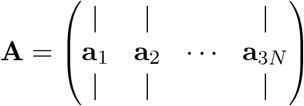

**Λ** is a 3*N* × 3*N* diagonal matrix with diagonal values equal to the associated normal modes’ squared angular frequencies *ω*^2^. The potential function used in this study is:

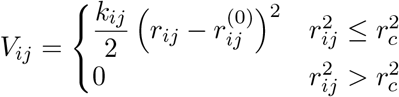

where *r_c_* is a cut-off radius, which for this work is set at 8Å; while *r*^(0)^ is the equilibrium distance between nodes derived form PDB crystallographic structure. For the wild-type protein, all spring constants are equal *k_ij_* = *k*= 1 kcal Å^−2^ mol^−1^.

#### Cross-correlation of Motion

The cross-correlation, *C*, is estimated between an ENM node pair as a normalised dot product sum between their normal mode eigenvectors over *v* modes.

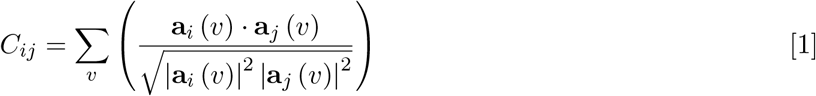

*C* value of 1 implies perfectly correlated motion, −1 perfectly anti-correlated motion and 0 implies totally non-correlated motion.

#### Normal Mode Fluctuation Free Energy

Using statistical mechanics it is possible to calculate an estimate to the fluctuation free energy of a system using the frequency of vibrations such as the normal modes. For this method, the partition function for the quantum harmonic oscillator (31), *Z*, for normal mode *k* is given as

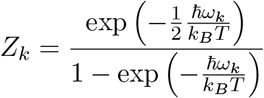

where *k_B_* is the Boltzmann’s constant, 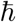 is the reduced Planck’s constant, *T* is temperature in Kelvin and *ω* is, already mentioned, angular frequency. Gibbs free energy (for a given mode) expressed in terms of partition function, with an approximation of little change in volume, can be written as

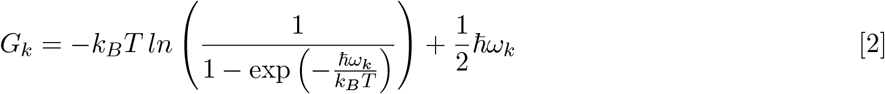

#### Ligand Dissociation Constant

When free energy change Δ*G* (**SI, Sec.**) is known for a dissociation reaction, corresponding dissociation constant *K* can be estimated via

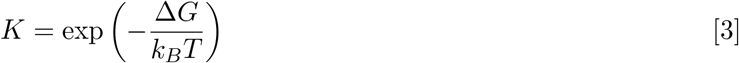

## Data and Code Availability

All data as well as the code to make the figures in this manuscript are available at https://github.com/burano/CompDynAlloMpro.

DDPT source code can be accessed at https://sourceforge.net/projects/durham-ddpt.

## ACKNOWLEDGMENTS

ID is grateful for computational support from the University of York high performance computing service, The Viking Cluster, especially Dr. Andrew Smith. TCBM acknowledges support from the EPSRC(UK) through an Established Career Fellowship in the Physics of Life programme. We are grateful to Dr. Alice von der Heydt, Prof. Peter O’Brian and Dr. Sarah Harris for useful discussions

## Footnotes

* From now on, M^pro^ will refer to the SARS-CoV-2 main protease protein

## Notes

https://github.com/burano/CompDynAlloMpro

### Competing Interest Statement

The authors have declared no competing interest.

