## Supplementary material for "Computational Analysis of Dynamic Allostery and Control in the SARS-CoV-2 Main Protease": SI

#### This PDF file includes:

Supplementary text
Figs. S1 to S5
SI References

### Supporting Information Text

**A. PDB structures.** Atoms of bound molecules were manually removed from the original 6LU7 PDB file to make up the three protein forms: ligand-free (apo); singly-bound ligand to chain A (holo1); each chain, A and B, posses one ligand (holo2).

**B. DDPT.** This section enumerates the routine used to generate and analyse $M^{pro}$ (6LU7) ENM via the Durham Dynamic Protein Toolbox (DDPT) (1). As an example, the routine is carried out explicitly here for the 6LU7 holo1 form.
First, the ENM interaction matrix is generated using the GENENMM function in DDPT:

GENENMM -pdb 6lu7_holo1.pdb -c 8 -het -fcust ffile.txt -mass -ca -res

The crystallographic structure of a protein is assigned by `-pdb` flag. The cut-off distance (`-c`, by default 12 Å) was set to 8 Å and heteroatoms (`HETATOM` in PDB notation) from the PDB file were included using `-het`, if required. Hookean spring strengths (by default 1 kcal $\text{Å}^{-2}$ $\text{mol}^{-1}$) around selected residues can be altered via the `-fcust` flag. The associated `ffile.txt` specifies which residue springs' moduli are modified. By default, all ENM nodes' masses (atoms from PDB structure are converted into ENM nodes) are set to the same fixed value, 1 amu, but actual atomic masses (`-mass`) were used in our ENM. The `-ca` flag creates an ENM model where amino acids' Cα atoms form nodes. In combination with `-ca`, the `-res` flag assigns the Cα nodes the whole residue mass. GENENMM Generates the interaction matrix for the ENM, written into `matrix.sdijf` file.
Next, the DIAGSTD function diagonalises the interaction matrix using small-block diagonalisation and iterative schemes (1).

DIAGSTD -i matrix.sdijf

where `-i` flag specifies input. The product of the diagonalisation is written into the `matrix.eigenfacs` file.
We calculate the inter-atom distance and cross-correlation of motion maps for the Cα ENM of the protein using SPACING and cross-cor functions, respectively.

SPACING -pdb 6lu7_holo1.pdb -ca -het

and

cross-cor -i matrix.eigenfacs -s 7 -e 31 -het

where `-s` and `-e` flags indicate the first and the last normal modes to include in calculation. Note, the first six normal mode frequencies, which correspond trivially to translational and rotational motion, are equal to zero.
Finally, fluctuation free energies for the set of the normal modes are calculated via the FREQEN function as follows

FREQEN -i matrix.eigenfacs -s 7 -e 106

The temperature ($T$) value used to calculate the partition function $Z$ and Gibbs free energy $G$ was 298 K, the default value for the FREQEN function.

***B.1. Fluctuation Free Energy Approximation.*** For convenience, DDPT estimates fluctuation free energy for each normal mode in dimensionless units $\frac{G}{k_BT}$ as follows

$$\frac{G}{k_BT} = -ln\left(\frac{1}{1-\exp\left(-\frac{\hbar\omega}{k_BT}\right)}\right) + \frac{1}{2}\frac{\hbar\omega}{k_BT}$$

Normal mode frequencies of global modes in proteins are in acoustic regime (less than 10 THz) (2, 3). This fact is especially true for the slowest modes we investigated in this study. In this classical limit, $\frac{\hbar\omega}{k_BT} \ll 1$ at 298 K temperature. Therefore, FREQEN function in DDPT employs the following expression for fluctuation free energy calculation

$$\frac{G}{k_BT} \approx -ln\left(\frac{1}{1-\exp\left(-\frac{\hbar\omega}{k_BT}\right)}\right) \qquad [1]$$

In the differences ($\Delta G$) and difference of a difference ($\Delta\Delta G$) in fluctuation free energy (**Eq. 2**) significance of $\frac{1}{2}\frac{\hbar\omega}{k_BT}$ term is negligible.

**C. Fluctuation Energy Convergence.** The allosteric free energy change is calculated using the fluctuation free energy (**Eq. 6**) change for three forms of the protein:

$$\begin{aligned}\Delta\Delta G &= \Delta G_2 - \Delta G_1\\ &= (G_{holo2} - G_{holo1}) - (G_{holo1} - G_{apo})\\ &= G_{holo2} - 2G_{holo1} + G_{apo}\end{aligned} \qquad [2]$$

For the apo form, the fluctuation free energy curve is smooth and stably converging with the number of modes included (**Fig. S1**). However, figure S2 shows a poorer convergence in the ratio of dissociation constants, due to the inherent noise-amplification in taking the difference of a difference in the (numerical) fluctuation free energies. However, the $K_2/K_1$ values fluctuate stably in the 1.00-1.02 range when the cut-off in mode number is between 25 and 35 modes. For this reason and three other stated in section 2 of the main paper text, we have chosen to stop summing at 25 non-trivial modes.
The 1-point mutational scan for the apo form is stable in respect of the identification of biologically active sites for increasing values of mode-sum cut-off (**Fig. S3A**). The same mutational scan, but for allosteric energy change, is less stable (for the same

reasons of numerical-difference as in the sensitivity of $K_2/K_1$ of figure S2 above) (**Fig. S3B**). Nevertheless, the identified bioactive sites persist stably with mode cut-off. As the sign of the stiffening modelling local mutation is changed, we observed a similar, and induced, sign change on in the allosteric response but, again, the patterns of bioactive residue persist. Visually inspection of the hydrophobic environment around residue 214 and 284-286 (**Fig. S4**) informed the choice of magnitudes of $k_R/k$ in our ENM that corresponds to the experimental mutation.
The identified candidate regions that exhibit allosteric activity in our ENM study (**Tab. 1**) were visualised in 3D space (**Fig. S5**).

**D. Definition of residues for dynamically allosteric control in SARS-CoV-2 M$^{pro}$ ENM.** We use results from the 1-point mutational apo scan (**Fig. 3A**) to suggest potential control residues on both homodimeric chains distant from the active sites. This map both captures significant fluctuation free energy change in the ENM dynamics of residue 214 and 284-286 mutations and the active residue patterns converge with fluctuation mode summation. In order to be considered a candidate control residue, the residue must pass two criteria:

1. Absolute fluctuation free energy change must be greater or equal to the smallest absolute value of experimentally evidenced dynamically allosteric residues (214 and 284-286) at $k_R/k$=0.25 or 4.00.

2. Cα node distance between the candidate residue and the catalytic active site residues (H41 and C145) in the ENM must be greater or equal to the double cut-off value (16 Å).

The first criterion filters out all residues which do not contribute to free energy change significantly; the second criterion ensures that the control effect of mutation or binding at the candidate residue is not caused by spatial proximity. We made only one exception (for E14) which is located on the homodimer interface; this showed desirable activity but was 15.1 Å away from C145 (**Fig. S5**). Residues that passed the two criteria above are documented in table 1.

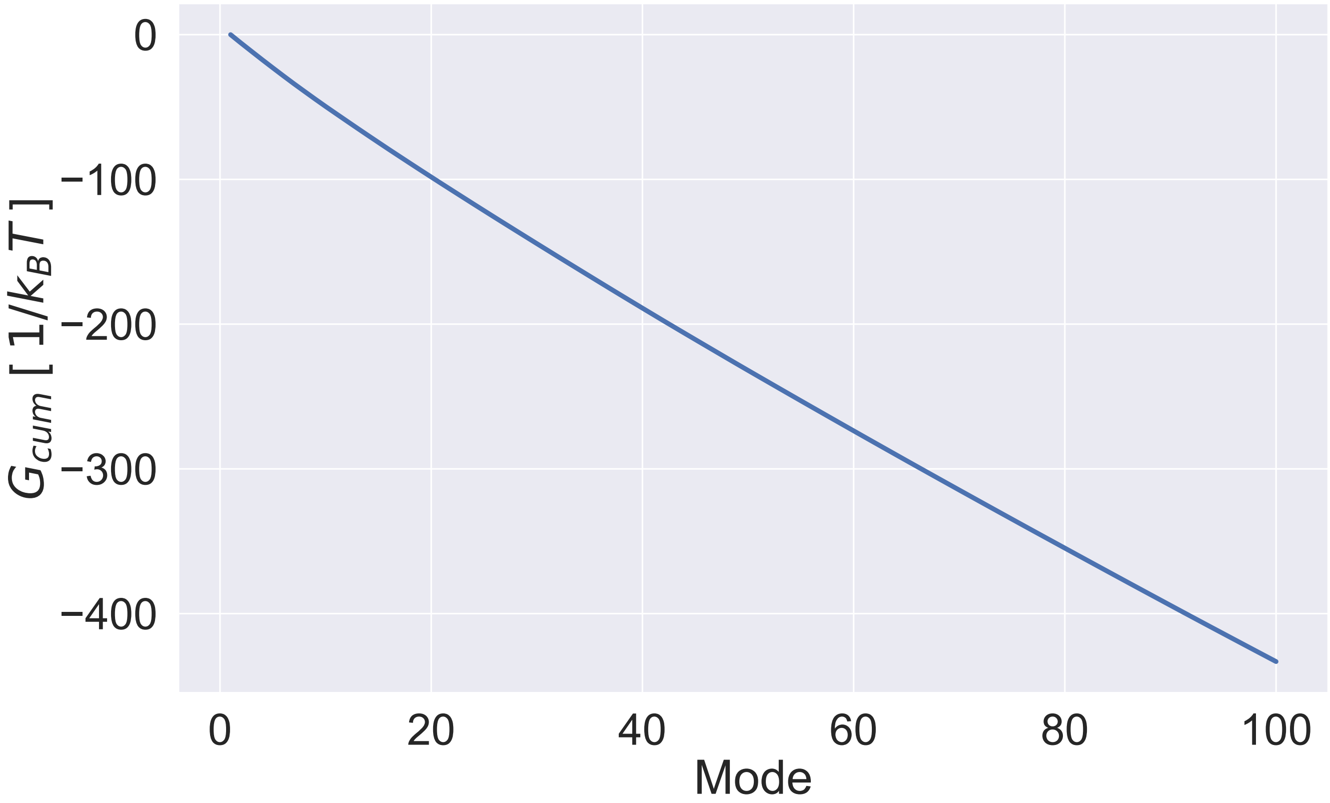


**Fig. S1.** Wild-type ligand-free SARS-CoV-2 $M^{pro}$ ENM fluctuation free energy dependence on mode summation at $298\,\mathrm{K}$. All ENM C$\alpha$ node masses correspond to amino acid mass, i.e. whole residue mass. Cut-off distance is 8 Å. All ENM spring constants are equal $1\,\mathrm{kcal}\,\mathrm{Å}^{-2}\,\mathrm{mol}^{-1}$

.

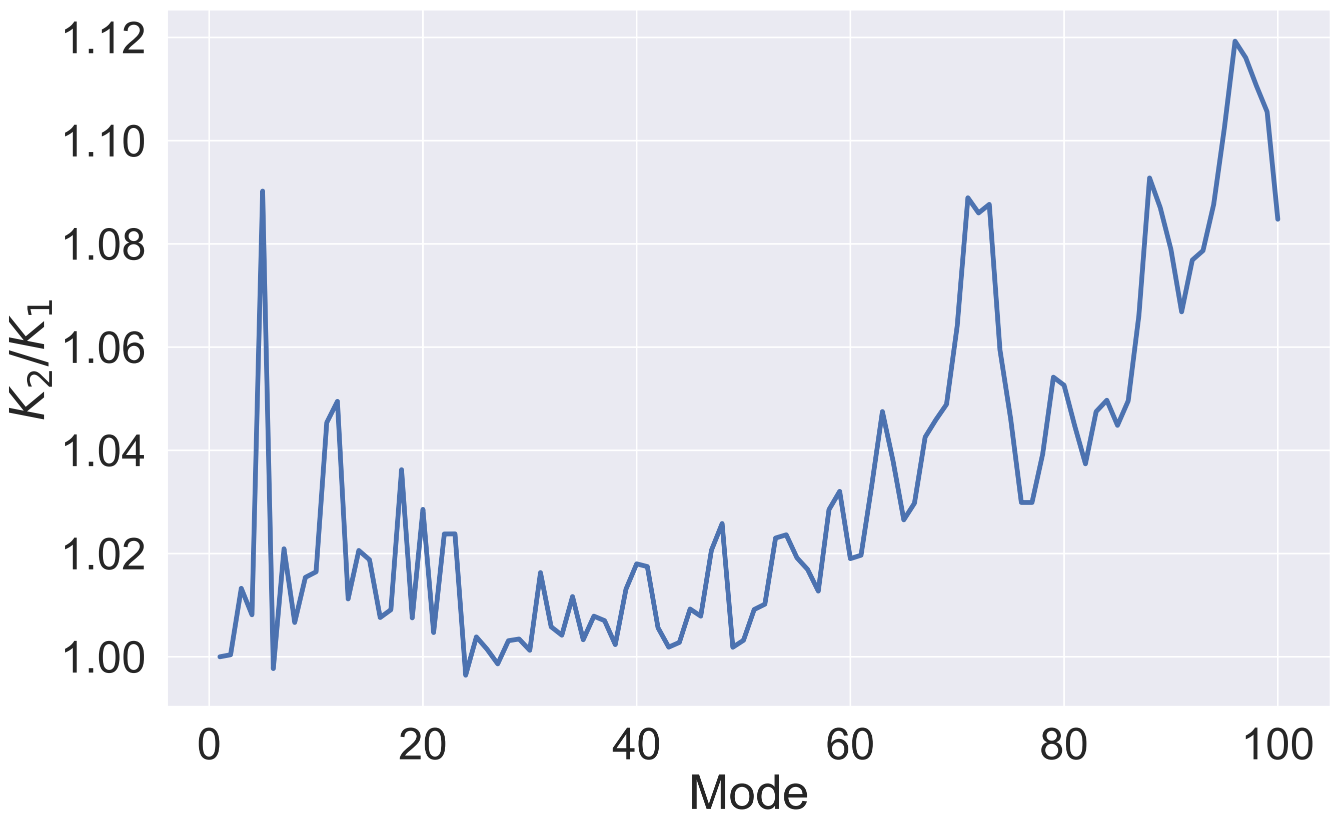


**Fig. S2.** Wild-type SARS-CoV-2 M$^{pro}$ ENM cooperativity dependence on mode summation. All ENM node masses correspond to amino acid mass a node been derived from; while ligand nodes mass is assigned based on the element. Cut-off distance is 8 Å. All ENM spring constants are equal $1\,\mathrm{kcal}\,\mathrm{Å}^{-2}\,\mathrm{mol}^{-1}$.

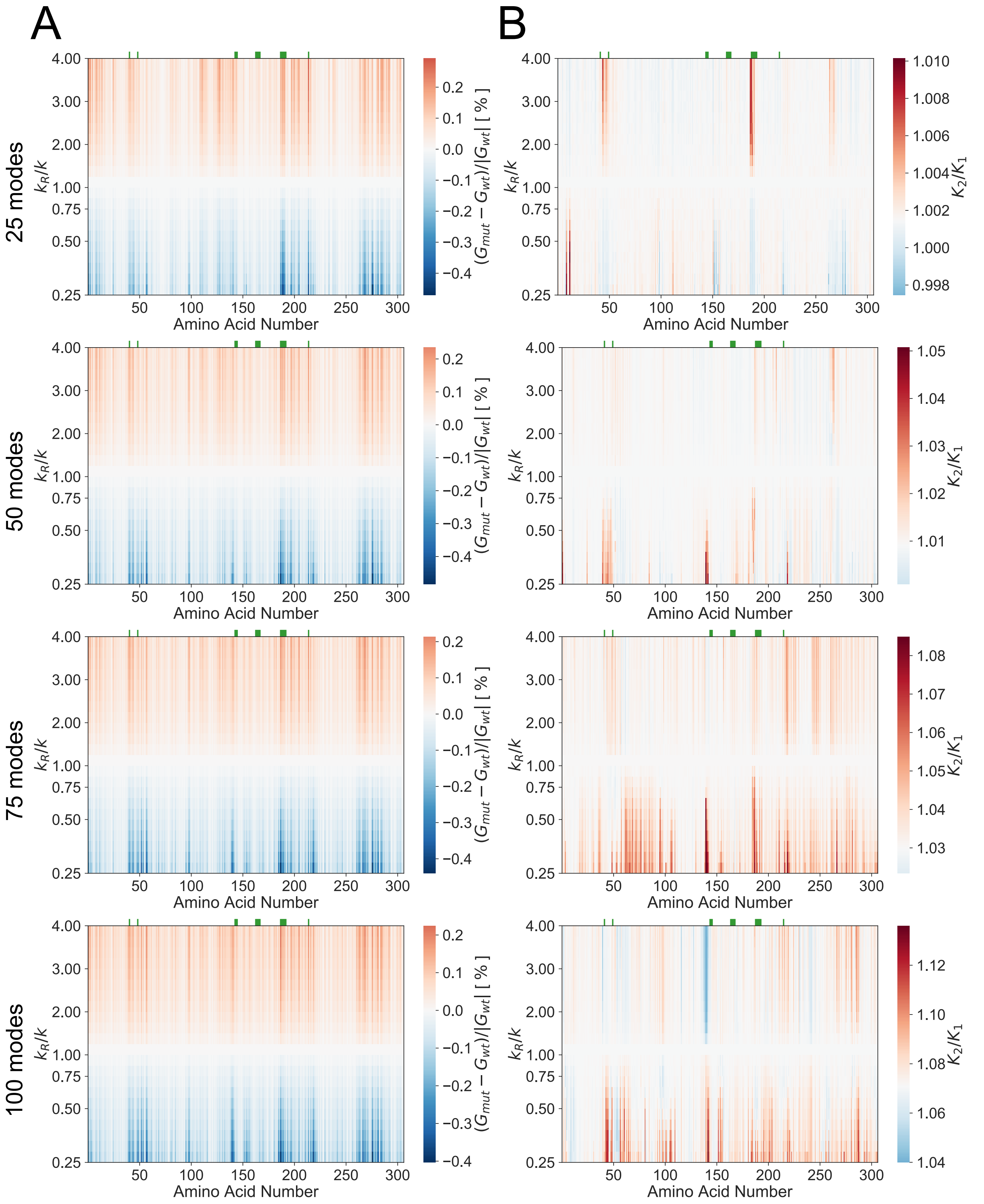


**Fig. S3.** 1-point mutational scan variation in $M^{pro}$ (6LU7) ENM over 4 increasing cumulative mode values: 25, 50, 75 and 100 modes. (A) 1-point mutational scan of apo ENM. (B) A map for the global control space of allostery in $M^{pro}$ calculated from the ENM.

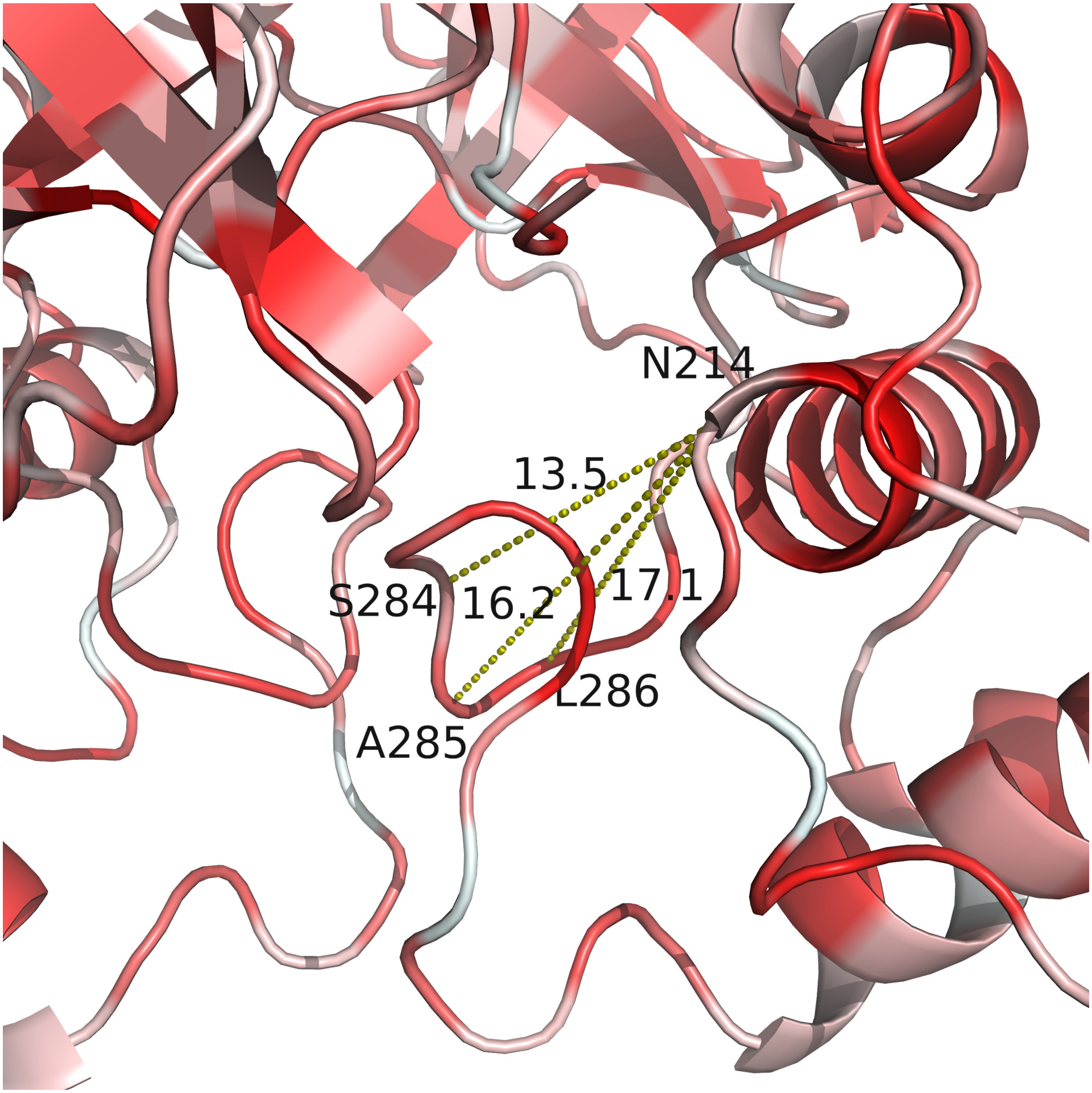


**Fig. S4.** Hydrophobic environment around residue 214 and 284-286 in SARS-CoV-2 $M^{pro}$ based on Eisenberg amino acid hydrophobicity scale (4). Red illustrates hydrophobic residues while white shows less hydrophobic (hydrophilic) residues. Euclidean distance between N214 and S284-A285-L286 is shown with yellow dashed lines in Å.

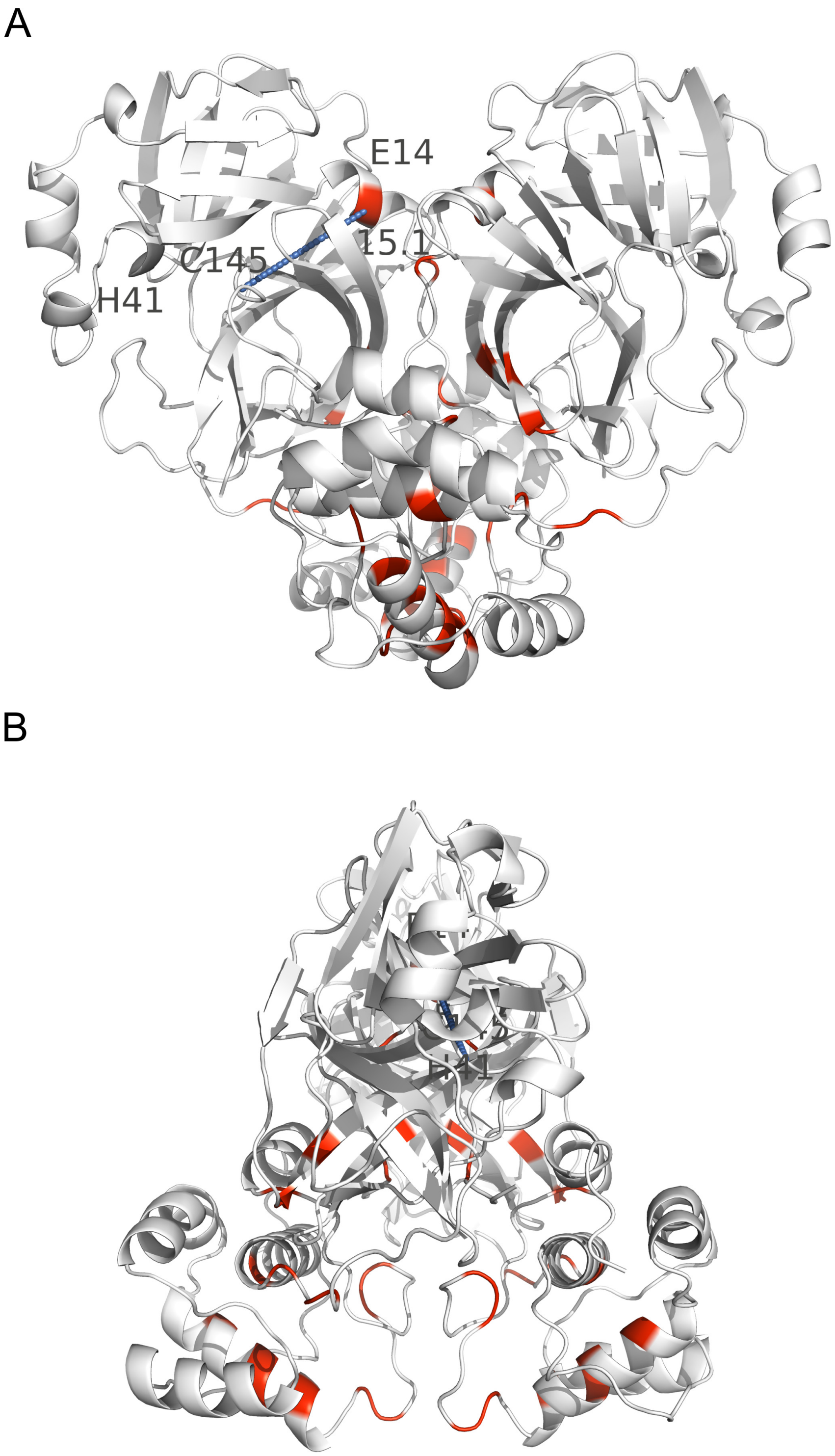


**Fig. S5.** Control residues candidates for dynamic allostery in $M^{pro}$, (A) front view and (B) side, that are distant from the catalytically active residues, H41 and C145. The identified residues are coloured in red on both homodimeric chains. The residues can be found in table 1 in the main manuscript text. A euclidean distance between E14 and C145 is shown with a blue dashed line and it equals $15.1\,\text{Å}$.
